# A DJ123 allele at the heading date quantitative trait locus *qHd7.1* promotes early heading without yield penalties under natural short-day and low fertility conditions in Madagascar

**DOI:** 10.1101/2024.10.23.619342

**Authors:** Katsuhiko Kondo, Kiyosumi Hori, Mbolatantely Rakotondramanana, Harisoa Nicole Ranaivo, Yoshiaki Ueda, Matthias Wissuwa

**Author notes:** **Corresponding author:** Matthias Wissuwa.

## Abstract

QTL analysis of heading date (Hd) was performed using a recombinant inbred line population derived from a cross between rice cultivars ʻIR64’ and ʻDJ123’. Phenotypic data was obtained from sites in Japan and Madagascar differing in photoperiod and soil fertility. The Japan site had long-days (LD) and high fertility, while the Madagascar site had short-days (SD) and low fertility conditions. Under LD in Japan, we discovered two Hd QTL on chromosome 7, *qHd7.1* and *qHd7.2*, whereas only one of these (*qHd7.1*) was detected under SD conditions in Madagascar. RILs carrying DJ123 alleles at both QTLs headed 9.5 days earlier in Japan (LD) compared to IR64 alleles, whereas the effect of DJ123 alleles at *qHd7.1* under SD conditions of Madagascar was 4 days and a combined effect of *qHd7.1* and *qHd7.2* did not exist. In Madagascar, early heading did not carry a yield penalty, whereas it caused reduced grain yield in Japan. These results suggest that RILs harboring the DJ123 allele of *qHd7.1* have high yield potential and good adaptation to low fertility conditions in Madagascar and that this can be achieved in a shortened cultivation period, improving resilience to effects of climate change for African rice farmers.

## Introduction

Rice is a staple food crop globally and improvement of rice yield is critical for ensuring food security for an increasing world population. To meet these food demands, it is not sufficient to solely improve yield potential of rice but adapting the rice crop to changing climatic conditions and cropping systems will be of importance. Heading date is a critical determinant of crop duration enabling crops to maximize the capture of daylight under specific ecological conditions, while also determining the ability for adaptation to new geographical locations and changing climatic conditions.

Rice is a traditional staple food in parts of West Africa and Madagascar, and it is increasingly becoming an important food in sub-Saharan Africa (SSA) due to population growth and a shift in consumer preference for rice (Balasubramanian *et al*. 2007). However, rice cultivation in SSA has many problems such as poor soil and water management. Declining soil fertility in SSA, especially phosphorus (P) deficiency, is a fundamental impediment to agricultural growth and a major reason for the slow growth in rice production (Balasubramanian *et al*. 2007, Tsujimoto *et al*. 2019). P deficiency typically delays maturity by one to two weeks, mainly due to delayed onset of tillering (Dobermann *et al*. 2000). Such increase in crop duration can affect rice yields negatively where water availability or cool temperatures are limiting factors toward the end of the growing season. Shortening the cultivation period, as well as increasing the yield potential, is critical to the demands of the African farmer.

Rice is a tropical short-day (SD) plant, and flowering is initiated as day length decreases below a critical threshold. Rice cultivars have diversified after domestication by adjusting their photoperiod sensitivity, allowing them to expand to a wide range of latitudes, from 45°N to 35°S (Izawa 2007, Monfreda *et al*. 2008). Rice has an evolutionarily conserved floral induction mechanism, which allows for the photoperiodic control of flowering. As a result, rice cultivar adaptation to regions with various daylength required selection for novel alleles that modify the original SD-specific photoperiod response to have flowering induced under long-day (LD) as well as short-day (SD) conditions.

So far, hundreds of rice QTLs affecting heading date have been identified by various genetic approaches and many of the underlying flowering time genes have been identified (Yamamoto *et al*., 2012, Hori *et al*., 2016, Yano *et al*., 2016, Yano *et al*., 1997, Yamamoto *et al*., 1998, Takahashi *et al*., 2001, Xue *et al*., 2008, Yan *et al*., 2011, 2013, Matsubara *et al*., 2012, Hori *et al*., 2013). Among these, several major QTLs have pleiotropic effects on heading date, plant height and grain yield.

Flowering time of rice is regulated by two independent floral pathways; one is the *Heading date 1* (*Hd1*)-dependent (*OsGI*-*Hd1*-*Hd3a*) pathway, the other being the *Early heading date* 1 (*Ehd1*)-dependent pathway. The *OsGI*-*Hd1*-*Hd3a* pathway is evolutionally conserved with the *GI-CO-FT* pathway in *Arabidopsis* (Shrestha *et al*. 2014). *OsGI* encodes an ortholog of *GIGANTEA* (*GI*), which is related to the circadian clock in *Arabidopsis* (Kim *et al*. 2007). *Hd1* is upregulated by OsGI and Hd1 activates the expression of *Hd3a* to promote rice heading under both SD and LD conditions (Hayama *et al*. 2003, Zhang *et al*. 2017). The *Ehd1*-*Hd3a* pathway of rice is not present in *Arabidopsis* and is regulated by genes upstream of *Ehd1. Ghd7, Ghd8, Ghd7.1/OsPRR31, Hd16, OsCOL4* and *OsCOL10* downregulate the expression of *Ehd1*, and cause late flowering under LD conditions (Xue *et al*. 2008, Lee *et al*. 2010, Yan *et al*. 2011, Hori *et al*. 2013, Yan *et al*. 2013, Tan *et al*. 2016).

*Grain number, plant height, and heading date 7* (*Ghd7*) is a rice-specific gene encoding a CCT domain protein, and it has effects on agronomic traits including number of grains per panicle, plant height and heading date. Enhanced expression of *Ghd7* delays heading and increases plant height and panicle size under LD conditions (Xue *et al*. 2008). *OsPRR37/Ghd7.1/Hd2* encodes a PSEUDO-RESPONSE REGULATOR (PRR) 7-like protein harboring the CCT domain. PRRs are essential circadian clock components in *Arabidopsis* (Alabadi *et al*. 2001, Farre and Kay 2007, Ito *et al*. 2009, Kaczorowski and Quail 2003, Nakamichi *et al*. 2007, Yamamoto *et al*. 2003). The rice orthologue to *Arabidopsis PRR7*, *OsPRR37*, represses heading under LD conditions and increases grain yield (Koo *et al*. 2013, Yan *et al*. 2013, Gao *et al*. 2014). Natural variations in *OsPRR37/Ghd7.1* also contribute to rice cultivation at a wide range of latitudes (Koo *et al*., 2013, Yan *et al*., 2013).

The early heading rice variety DJ123 maintains high P uptake from P-deficient soils and carries a QTL for high root efficiency on chromosome 11 (Mori *et al*. 2016). It further carries a QTL for P utilization efficiency at a different location on chromosome 11 (Wissuwa *et al*. 2015). IR64 is a high-yielding irrigated rice variety developed by the International Rice Research Institute (IRRI) that is widely grown in Asia (Mackill and Khush 2018). Therefore, the combination of high yield potential in IR64 with early heading and high P efficiency in DJ123 could lead to the development of promising rice varieties that have improved adaptation to low soil fertility or P-deficient soil conditions in Africa.

To this end we have crossed donor DJ123 with IR64 and developed a population of recombinant inbred lines (RILs) that was phenotyped in Japan and Madagascar under contrasting conditions with regard to day length and soil fertility. The objectives of this study were (1) to conduct QTL analysis for heading date in the RIL population under LD and high soil fertility in Japan, and under SD and low fertility in Madagascar; and (2) to further clarify whether RILs can be identified that combine early heading with high yield potential.

## Materials and Methods

### Plant materials and field experiment

We used IR64 (*indica*) and DJ123 (*aus*) as parental varieties in this study and following their cross recombinant inbred lines (RILs) were derived by single-seed descent (SSD) method as reported previously (Ueda *et al*. 2024). The QTL mapping population consisted of 288 RILs that were evaluated under irrigated lowland conditions at two locations in the F_4_ and F_5_, respectively. The first trial was conducted with F_4_ RILs from April to September 2017 at the Japan International Research Center for Agricultural Sciences (JIRCAS) in Tsukuba, Japan, (36°05’ N, 140 °08’E), while the second trial using derived F_5_ RILs was conducted from November 2017 to May 2018 at a farmer’s field in Ankazo, Madagascar, (19°40’ S, 46 °33’E). The plants were grown at day length greater than 13.5 hours in Japan (considered long-day (LD) conditions), and at average day length of less than 12.5 hours in Madagascar (natural short-day (SD) conditions) (Supplemental Fig. 1). The median of total P concentration in soils is 1.43g P kg^-1^ in Japan, while it is 0.68g P kg^-1^ in Madagascar (Yanai, Okada, and Yamada 2012, Nishigaki *et al*. 2019). Using 28-day-old seedlings that had been raised in a seedling nursery, single plants were transplanted into the field using a distance of 20 cm between plants within a row and 30 cm between rows. Each line had thirteen plants transplanted per row. Five individual plants were selected from the middle of each line for data collection of five traits: plant height (PH), days to heading (DTH), panicle dry weight (PDW), panicle number per plant (PN) and shoot dry weight (SDW). Observations for these traits were averaged over five plants.

### QTL analysis

The 288 F_4_ RILs were genotyped using the <u>K</u>ompetitive <u>A</u>llele <u>S</u>pecific <u>P</u>olymerase chain reaction (KASP) assay (Semagn *et al*. 2014) by LGC Genomics (Hoddesdon, Hertfordshire, UK) (https://www.lgcgroup.com/products/kasp-genotyping-chemistry/). 185 polymorphic KASP markers were selected from the set of markers described by Pariasca-Tanaka *et al*. (2014) and the construction of the linkage map and QTL analysis were performed using the package QTL in R (R Statistical Computing, Broman *et al*. 2003). Genetic distances were estimated using the Kosambi mapping function (Kosambi 1944), and putative QTLs were detected by composite interval mapping (CIM). CIM thresholds at the 5% level of significance were derived from 1,000 times of permutation test (Churchill and Doerge 1994). The additive effect and the percentage of phenotypic variance (*R*^2^) were estimated at the peak of LOD scores.

### Sequence analysis of Ghd7 and OsPRR37

To analyze the sequence of *Ghd7* and *OsPRR37*, PCR products were amplified using PrimeSATR GXL DNA Polymerase (TaKaRa, Ohtsu, Japan) from genomic DNA of IR64 and DJ123. Gene specific primers were used for *Ghd7* and *OsPRR37* (Supplemental Table. 1). Amplified DNA fragments were purified and sequenced by ABI3130xl sequencers (Applied Biosystems, Foster City, CA, USA). The promotor sequence of *Ghd7* were obtained from *de novo* assembly of next generation sequence data by the Cold Spring Harbor Laboratory (Schatz *et al*. 2014). Assembly of the sequence data and alignment of the amino acid sequences were generated using the ATGC program and GENETYX (Genetyx Corp., Tokyo, Japan). Phosphorylation sites of Ghd7 were predicted using the NetPhos 3.1 server (http://www.cbs.dtu.dk/services/NetPhos/) (Blom *et al*. 1999).

## Results

### Phenotypic differences between parents IR64 and DJ123 and variation within RILs

All traits investigated in this study exhibited significant differences between IR64 and DJ123 in the two different field conditions. DJ123 reached the heading stage earlier than IR64 in both Japan and Madagascar; in Japan DTH for DJ123 was 88 days compared to 114 days for IR64 (26 days difference) (Fig. 1A), whereas in Madagascar DTH for DJ123 was 98 days compared to 107 days for IR64 (9 days difference). DJ123 was significantly taller than IR64 and this difference was more pronounced in Madagascar (+26 cm) than in Japan (+20 cm) (Fig. 1B). Plant height (PH) in Madagascar was reduced relative to Japan, indicating that growth conditions were more favorable at the Japanese site. That was confirmed by reduced panicle number (PN) and panicle dry weight (PDW) in Madagascar (Fig. 1C, D). IR64 produced significantly more panicles and more PDW than DJ123 at both sites.

**Fig. 1.**
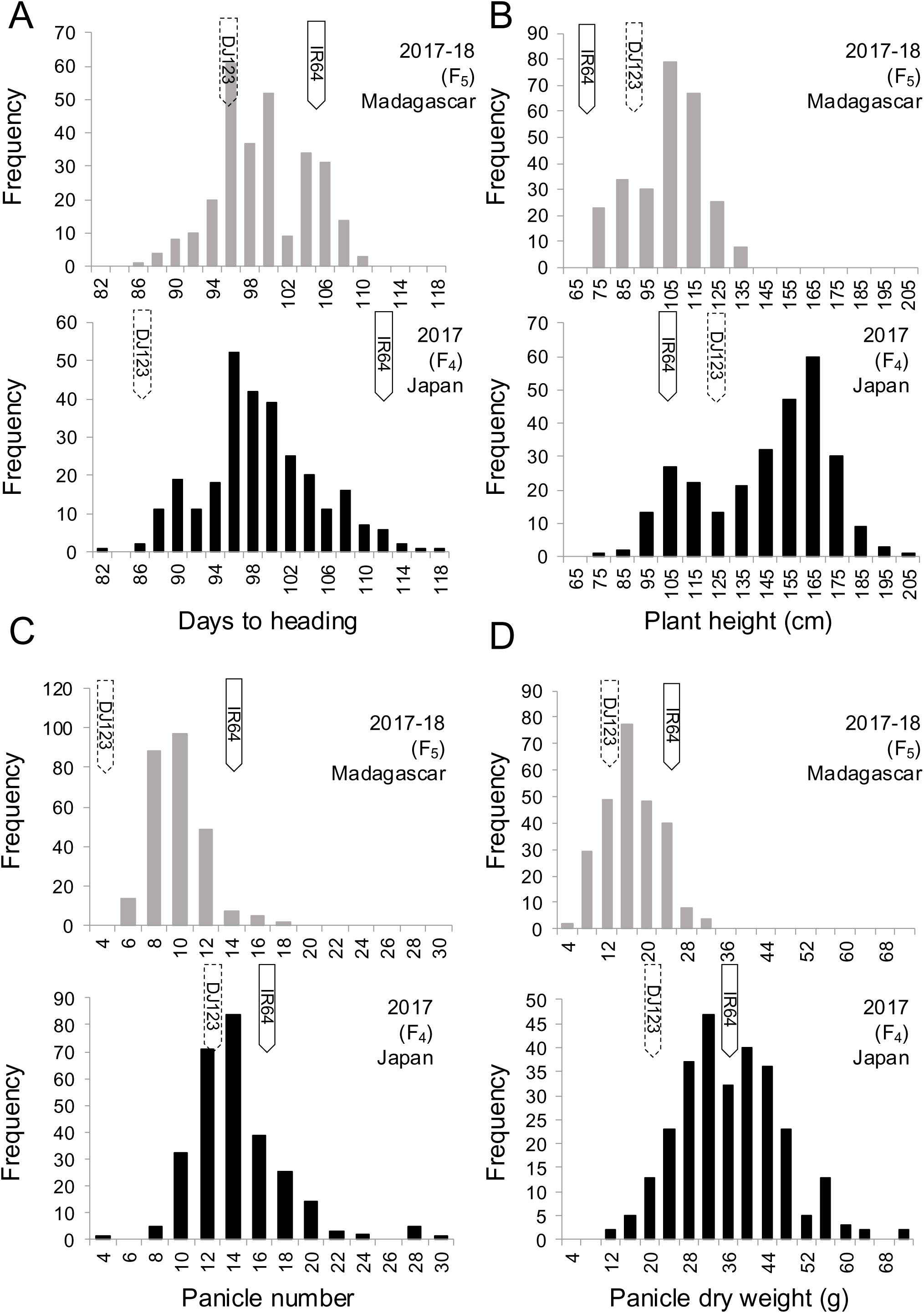
Frequency distribution of agronomical traits obtained in Japan and Madagascar. Data for days to heading (A), plant height (B), panicle number (C) and panicle dry weight (D) obtained from RILs derived from a cross between IR64 and DJ123 are shown. Gray and black bars indicate phenotypic values obtained in Madagascar and Japan, respectively. Solid and dotted line arrows indicate average values for the parents (IR64 and DJ123).

All traits showed a continuous distribution and transgressive segregation in RILs. Mean values of PH, PN, and PDW among RILs in Madagascar were significantly lower than in Japan, again confirming less fertile soil conditions in Madagascar.

### QTL analysis for DTH under two field conditions

We performed QTL analysis on DTH obtained from the two locations differing in latitude and soil conditions. Two major QTL for DTH, *qHd7.1* and *qHd7.2*, were detected in Japan, while only one of these major QTL (*qHd7.1*) was identified in Madagascar (Fig. 2; Table 1). Under LD conditions in Japan, *qHd7.1* and *qHd7.2* explained 13.7% and 16.2% of the phenotypic variance observed (Table 1). Under SD conditions in Madagascar, *qHd7.1* explained 17.7% and 17.5% of the phenotypic variance in the two replicate field plots (Table 1). The DJ123 allele at *qHd7.1* accelerated heading in both SD and LD conditions, while *qHd7.2* on chromosome 7 affected DTH only under LD conditions.

**Fig. 2.**
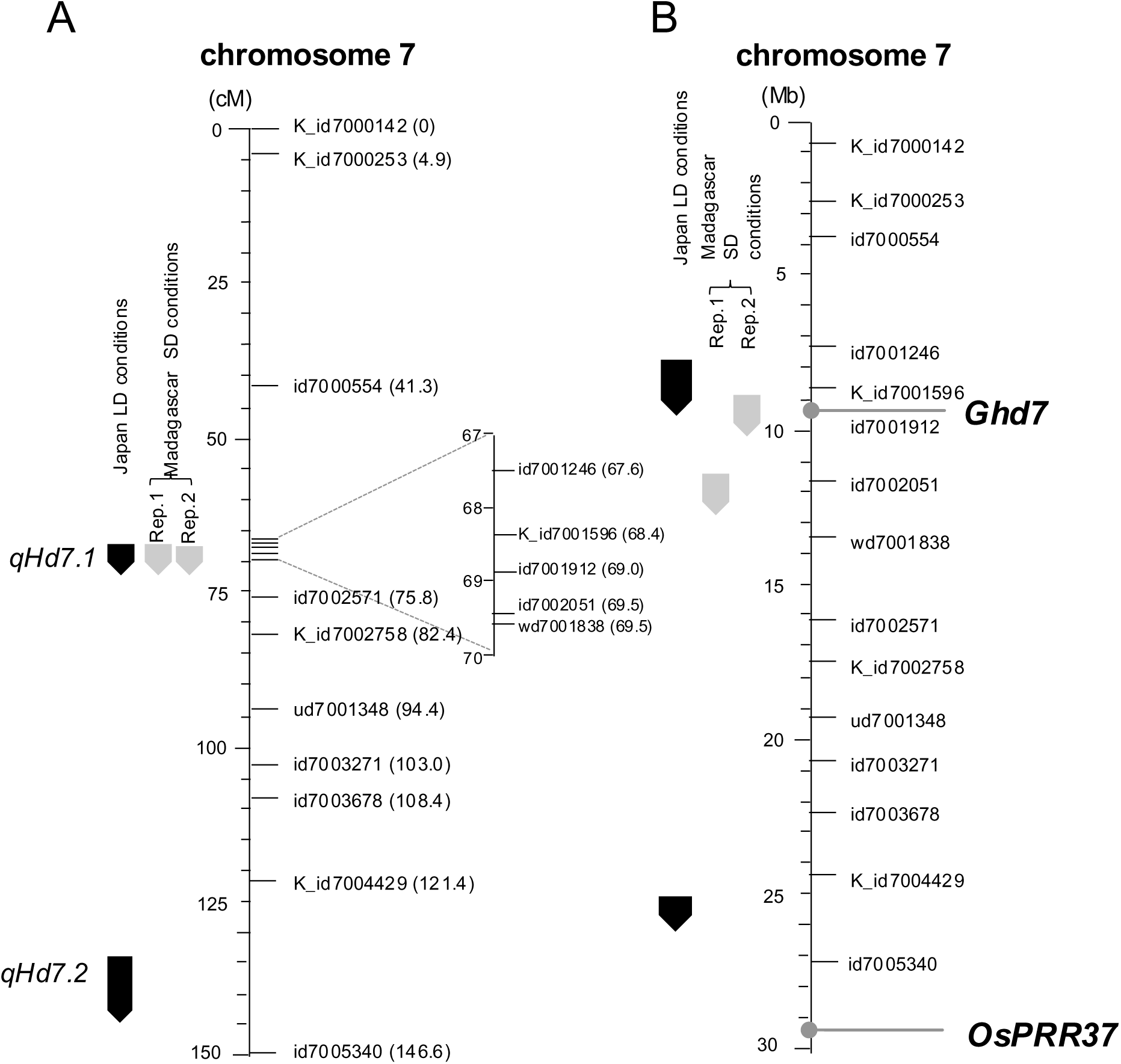
Genetic and physical maps of heading date QTLs detected on chromosome 7. (A) Linkage map developed using RILs from a cross between IR64 and DJ123. Distances are shown in centimorgans. (B) Physical map of the two QTLs and physical distances based on the reference Nipponbare genome. Black and gray arrows indicate detected QTLs from Japan LD or Madagascar SD conditions, respectively.

**Table 1.** Putative QTLs for days to heading detected in Japan and Madagascar using a RIL population derived from a cross of IR64 × DJ123.

| Genotype/<br>phenotype | Location | QTL | Chr. | Flanking<br>marker | LOD | Additive<br>effect <sup>a</sup> | R<sup>2</sup> <sup>b</sup> |
| --- | --- | --- | --- | --- | --- | --- | --- |
| F<sub>4</sub>/F<sub>4</sub> | Japan | *qHd7.1* | 7 | K_id7001596 | 11.5 | -2.61 | 13.7 |
|  |  | *qHd7.2* | 7 | K_id7004429<br>id7005340 | 13.4 | -2.73 | 16.2 |
| F<sub>4</sub>/F<sub>5</sub> | Madagascar<br>Rep.1 | *qHd7.1* | 7 | id7002051<br>wd7001838 | 11.5 | -2.33 | 17.7 |
| F<sub>4</sub>/F<sub>5</sub> | Madagascar<br>Rep.2 | *qHd7.1* | 7 | K_id7001596<br>id7001912 | 11.3 | -1.92 | 17.5 |

<sup>a</sup> Additive effect of the DJ123 allele compared with the IR64 allele.
<sup>b</sup> Percentage phenotypic variance explained by each QTL.

To further evaluate QTL effects, we compared the DTH between F_4_ or F_5_ lines homozygous for the IR64 or DJ123 allele at the two QTL regions (*qHd7.1* and/or *qHd7.2*). Individual plants homozygous for the DJ123 allele at *qHd7.1* accelerated heading by four days both in Japan and Madagascar (Fig. 3A). Plants homozygous for the DJ123 allele at *qHd7.2* were observed to head 6 days earlier in Japan but only 2 days earlier in Madagascar, compared to plants with the IR64 allele at *qHd7.2* (Fig. 3B). In addition, the combination of both *qHd7.1* and *qHd7.2* loci being homozygote for the *DJ123* allele was confirmed to accelerate DTH by 9.5 days under LD conditions in Japan but only by 3.5 days in Madagascar (Fig. 3C).

**Fig. 3.**
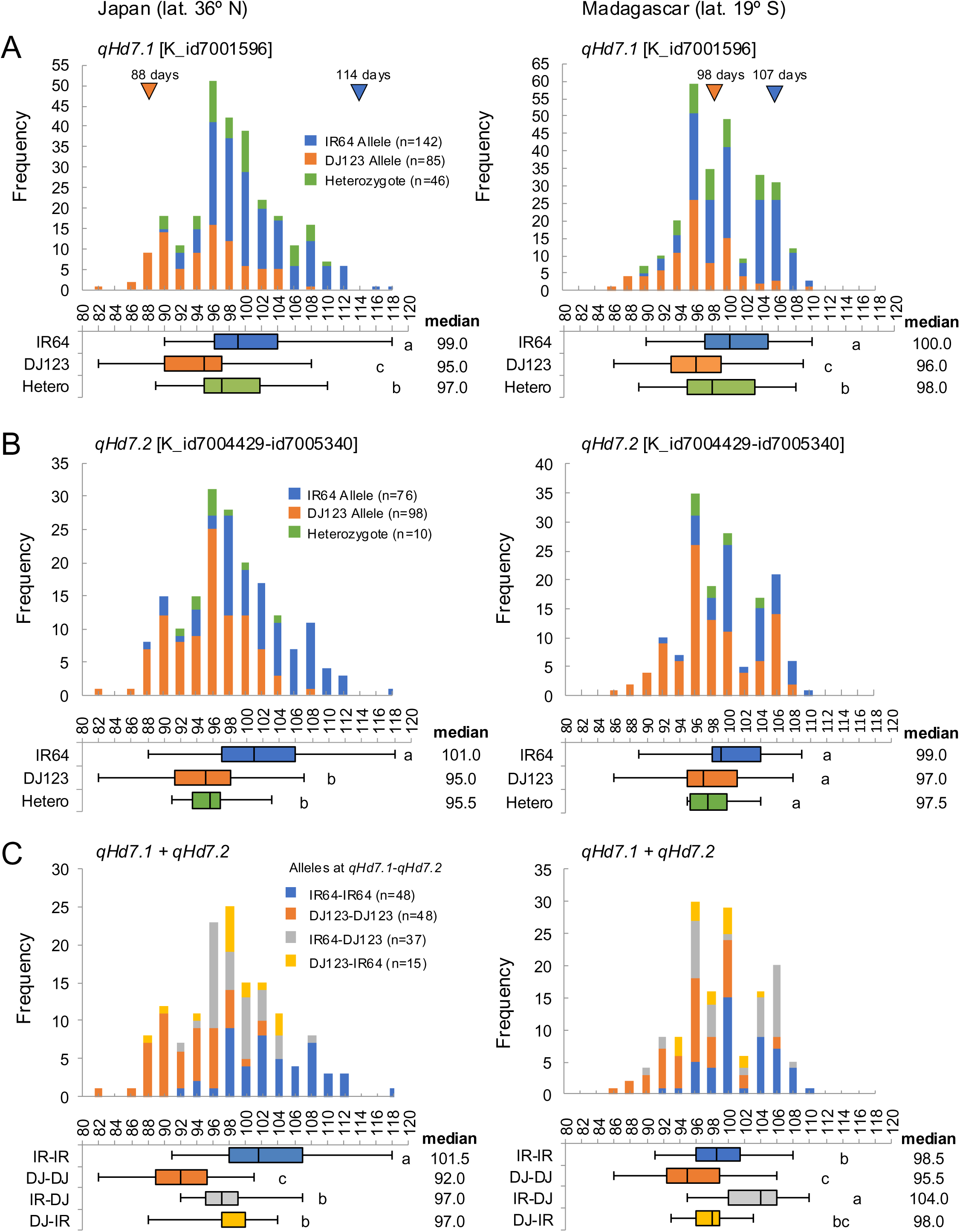
Frequency distribution of heading date in groups that differ for the alleles at *qHd7.1* and *qHd7.2*. Progenies of the RIL population were classified according to their allelic state at *qHd7.1* (A), *qHd7.2* (B) and both of *qHd7.1* and *qHd7.2* (C), and the distribution of their heading date is shown with histograms and box plots. Blue and orange arrows indicate the mean DTH of parents (IR64 and DJ123). Different alphabets indicate significant differences determined by two-sided Student’s *t*-test (*P* < 0.05). Fields in Japan were under LD conditions and supplied with NPK fertilizers, while those in Madagascar were under SD conditions with no fertilizer applied.

### Associations between heading date and grain yield under different field conditions

Over the entire RIL population, correlations between DTH, PDW, SDW and PH were not significant (Supplemental Fig. 2; Table 2). However, if the population was classified into groups with different allelic states at *qHd7.1* (+ *qHd7.2*), contrasting correlations between heading and PDW were detected between the groups, and this differed between LD and SD conditions. In lines homozygous for the DJ123 allele at *qHd7.1* (Fig. 4A) or at both loci (Fig. 4B), late heading was associated with higher PDW in LD Japan, but in Madagascar the opposite trend was observed as PDW tended to be higher in early heading lines. For lines with the IR64 allele, no trend was observed in Madagascar, while late heading reduced PDW in Japan (Fig. 4 A, B).

**Table 2.** Correlation coefficients among traits in the F_4_ and F_5_ lines derived from a cross of IR64 × DJ123.

| Japan (F<sub>4</sub>) |  |  |  |  | Madagascar (F<sub>5</sub>) |  |  |  |  |
| --- | --- | --- | --- | --- | --- | --- | --- | --- | --- |
|  | DTH | PDW | SDW | PH |  | DTH | PDW | SDW | PH |
| PDW | -0.02 |  |  |  | PDW | -0.09 |  |  |  |
| SDW | 0.00 | 0.45** |  |  | SDW | 0.19** | 0.45** |  |  |
| PH | 0.06 | 0.34** | 0.59** |  | PH | 0.09 | 0.12* | 0.42** |  |
| PN | -0.30** | 0.07 | 0.24** | -0.30** | PN | -0.01 | 0.20** | 0.23** | -0.29** |

Pearson's correlation coefficients among different traits are indicated. DTH, days to heading; PDW, panicle dry weight; SDW, shoot dry weight; PH, plant height; PN, panicle number. \* and \*\* indicate significance at the level of $P < 0.05$ and $P < 0.01$ , respectively.

**Fig. 4.**
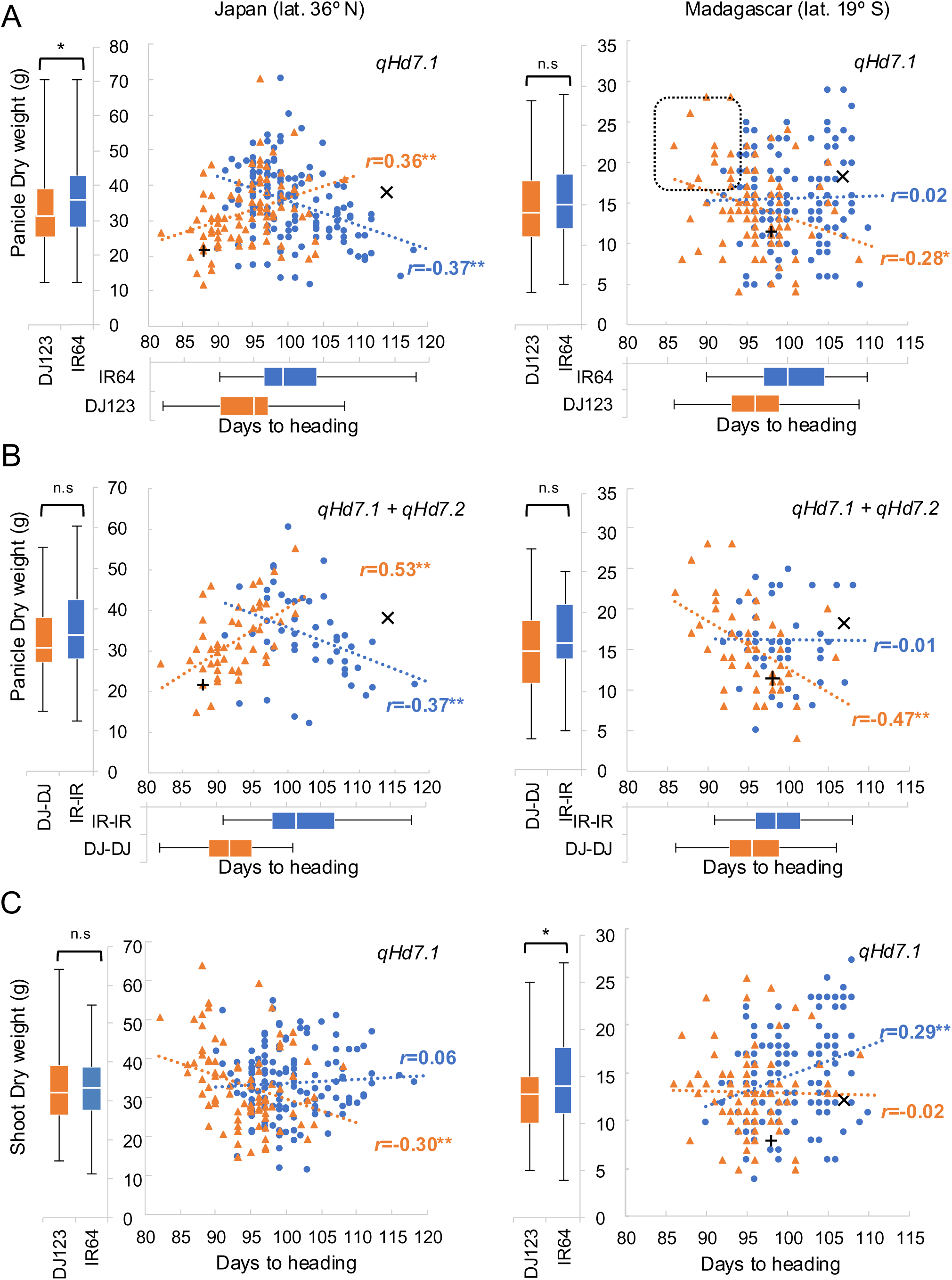
Correlation analyses between heading date and panicle traits. Scatter plots showing correlations between days to heading and panicle dry weight (A, B) and shoot dry weight (C) are shown. RIL population was classified according to their allelic state at *qHd7.1* (A, C) and both *qHd7.1* and *qHd7.2* (B). Blue and orange lines indicate the heading dates from IR64 and DJ123 homozygote alleles for *qHd7.1* (A, C) or the combination of *qHd7.1* and *qHd7.2*, respectively. × and + indicate the average of parents IR64 and DJ123, respectively. The dotted line box (the right panel of A) indicates the promising breeding lines under SD conditions in Madagascar. The asterisks (*) indicate significant correlation at the level of *P* < 0.05.

Given that the overall correlation between DTH and PDW was not significant, one point of interest is whether lines can be identified that combine earliness with high PDW. In Japan, earliest lines had lower yield despite having higher shoot biomass (Fig. 4C). In contrast, some early lines showed high PDW in Madagascar (Figs. 4, 5). One of the promising lines (DIN-129) was observed to head 8 days earlier than DJ123 (12 days earlier than IR64) under natural SD conditions in Madagascar, while PDW was 1.5 times larger than in IR64 (Fig. 5).

**Fig. 5.**
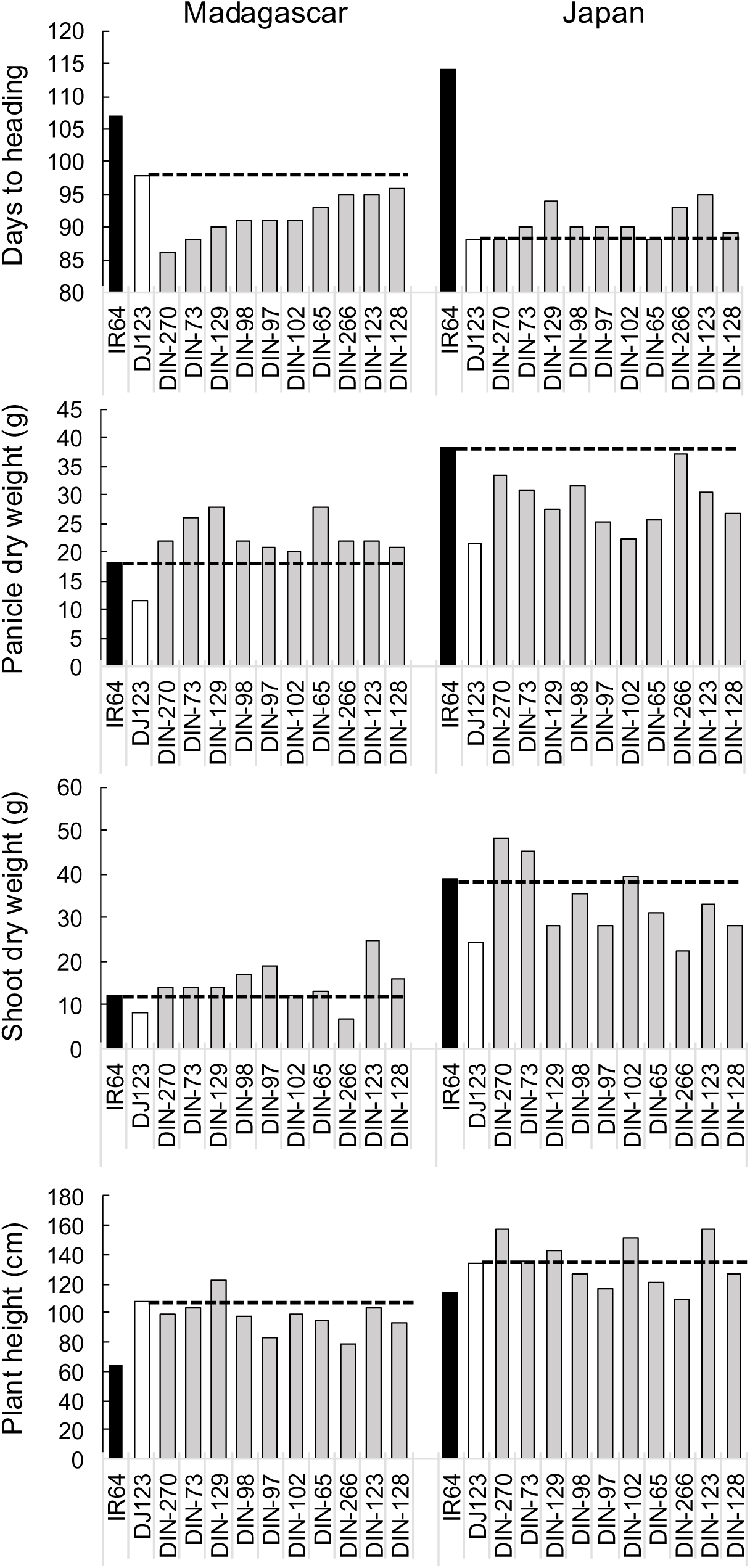
Agronomic traits of promising breeding lines from the RIL population in comparison to both parents. Phenotypic data from selected RILs and parents are shown.

### Comparison of amino acid sequences for candidate genes of qHd7.1 and qHd7.2

The two QTL (*qHd7.1* and *qHd7.2*) detected in this study are closely located to the previously reported Hd-related genes, *Ghd7/Hd4* and *OsPRR37/Ghd7.1/Hd2*, respectively (Fig. 2). We obtained the genomic and promoter sequences of *Ghd7* and *OsPRR3* from parents IR64 and DJ123 (Fig. 6). For the *Ghd7* gene, four amino acid differences were observed between IR64 and DJ123 alleles, with DJ123 being identical to the Nipponbare reference allele (Fig. 6A). Interestingly, the change to a serine residue at position 136 of Ghd7 was predicted to have added a new phosphorylation site in the IR64 allele. The analysis of the *Ghd7* promoter region furthermore showed that Nipponbare and DJ123 were nearly identical except for two SNPs, whereas 37 SNPs and one indel were detected in IR64 relative to the Nipponbare allele (Fig. 6B).

**Fig. 6.**
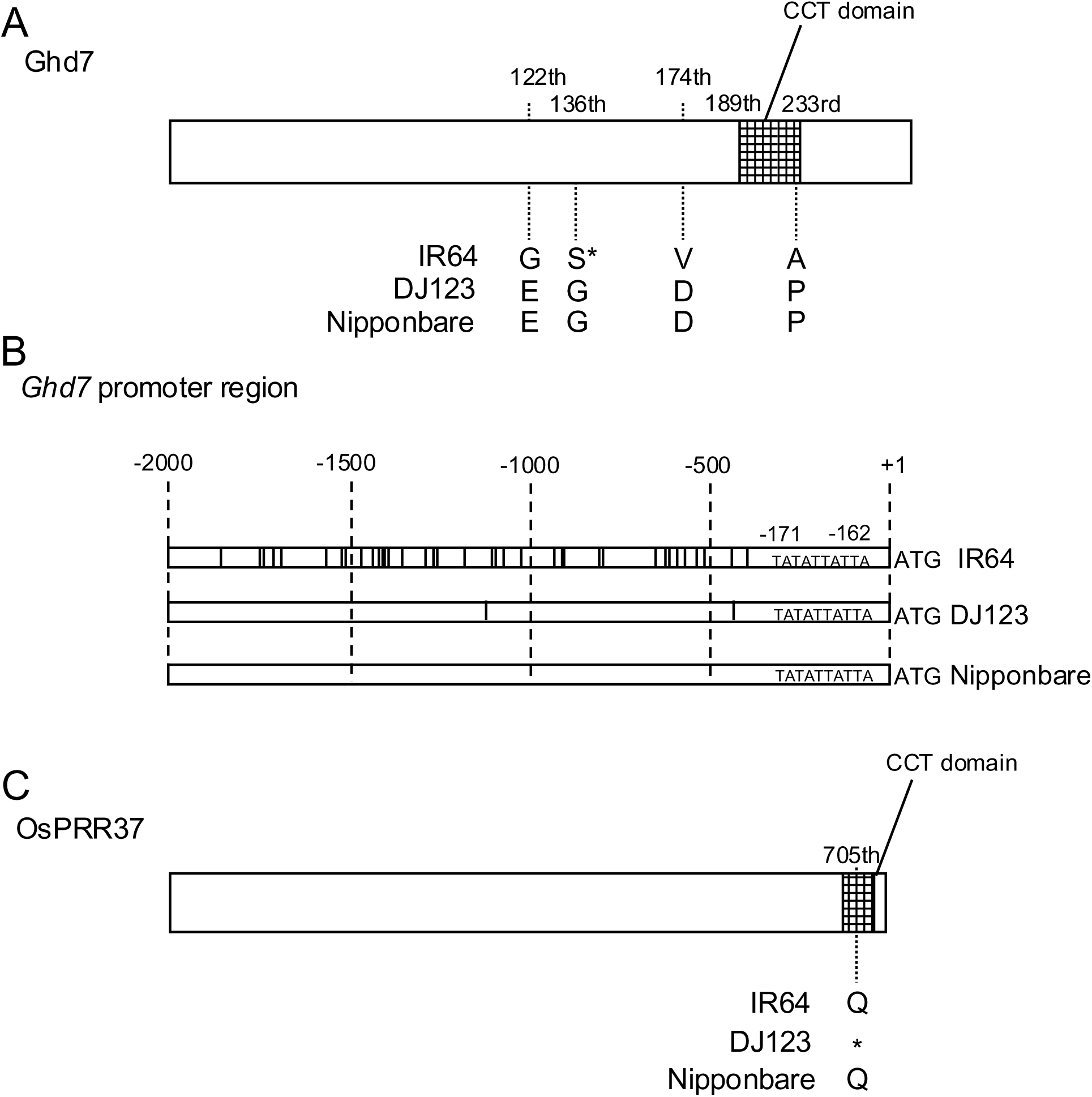
Amino acid and nucleotide sequences of *Ghd7* and *OsPRR37* in IR64, DJ123 and Nipponbare. (A) Predicted protein sequences of Ghd7. The phosphorylation site predicted by NetPhos 3.1 is indicated by an asterisk while the mesh shows the CCT domain. (B) Schematic diagram of the *Ghd7* promoter region. Differences with the Nipponbare sequence is indicated by vertical lines. (C) Predicted protein sequences of OsPRR37 with CCT domain indicated by a mesh.

For OsPRR37, the candidate gene at *qHd7.2*, the reverse situation was detected with IR64 being identical to the Nipponbare reference allele (Fig. 6C; Supplemental Fig. 4). One base pair substitution (compared with the Nipponbare allele) was detected in DJ123 and this substitution occurred in the conserved CCT motif, which is likely involved in nuclear localization. The base substitution at position 2113 (C-2133-T) caused a premature termination of translation in the CCT motif (Q-705-Stop) of the DJ123 allele (Fig. 6C; Supplemental Fig. 4).

## Discussion

### DJ123 alleles at qHd7.1 and qHd7.2 do not cause late heading under both SD and LD conditions

In this study, we detected two QTLs, *qHd7.1* and *qHd7.2*, for DTH from the Japan data. Only one QTL, *qHd7.1*, was detected in Madagascar. DJ123 alleles at each of these QTLs were able to accelerate DTH in both Japan and Madagascar. This result indicates that *qHd7.1* promotes early heading under both SD and LD conditions, but *qHd7.2* is not effective under SD conditions. These two QTLs (*qHd7.1* and *qHd7.2*) are closely linked to the previously reported *Ghd7* and *OsPRR37/Ghd7.1/Hd2*, respectively (Fig. 2). *Ghd7* and *OsPRR37* repress expression of *Ehd1*, resulting in late flowering under LD conditions. (Hori *et al*. 2016, Lee *et al*. 2010, Tan *et al*. 2016, Xue *et al*. 2008, Yan *et al*. 2013, Yan *et al*. 2011).

We obtained sequences of parents IR64 and DJ123 and compared their genomic and promoter regions of *Ghd7* and *OsPRR37*. In *OsPRR37*, the nucleotide sequence from DJ123 has a base substitution at position 2133 (C-2133-T), resulting in a premature termination of translation in the CCT motif (Q-705-Stop) (Fig. 6). This alteration of the CCT motif in the OsPRR37 was previously reported as a natural variation that causes early flowering under natural LD conditions (Murakami *et al*. 2005, Koo *et al*. 2013). This is consistent with our result linking *OsPRR37* closely with *qHd7.2,* for which the DJ123 allele caused earlier heading under LD conditions in Japan compared to the IR64 allele (Fig. 3). However, *qHD7.2* was not detected under the SD conditions in Madagascar (Fig. 2; Table 1), suggesting that *OsPRR37* might not be involved in photoperiodic regulation under SD conditions in the genetic background studied here (Fig. 7). Recently, it was reported that *OsPRR37* alternatively functions as an activator or a suppressor under natural SD conditions depending on the status of three genes (*Ghd7*, *Ghd8*, *Hd1*) (Zhang *et al*. 2019). The mechanism of regulation of *OsPRR37* expression under SD conditions remains unclear.

**Fig. 7.**
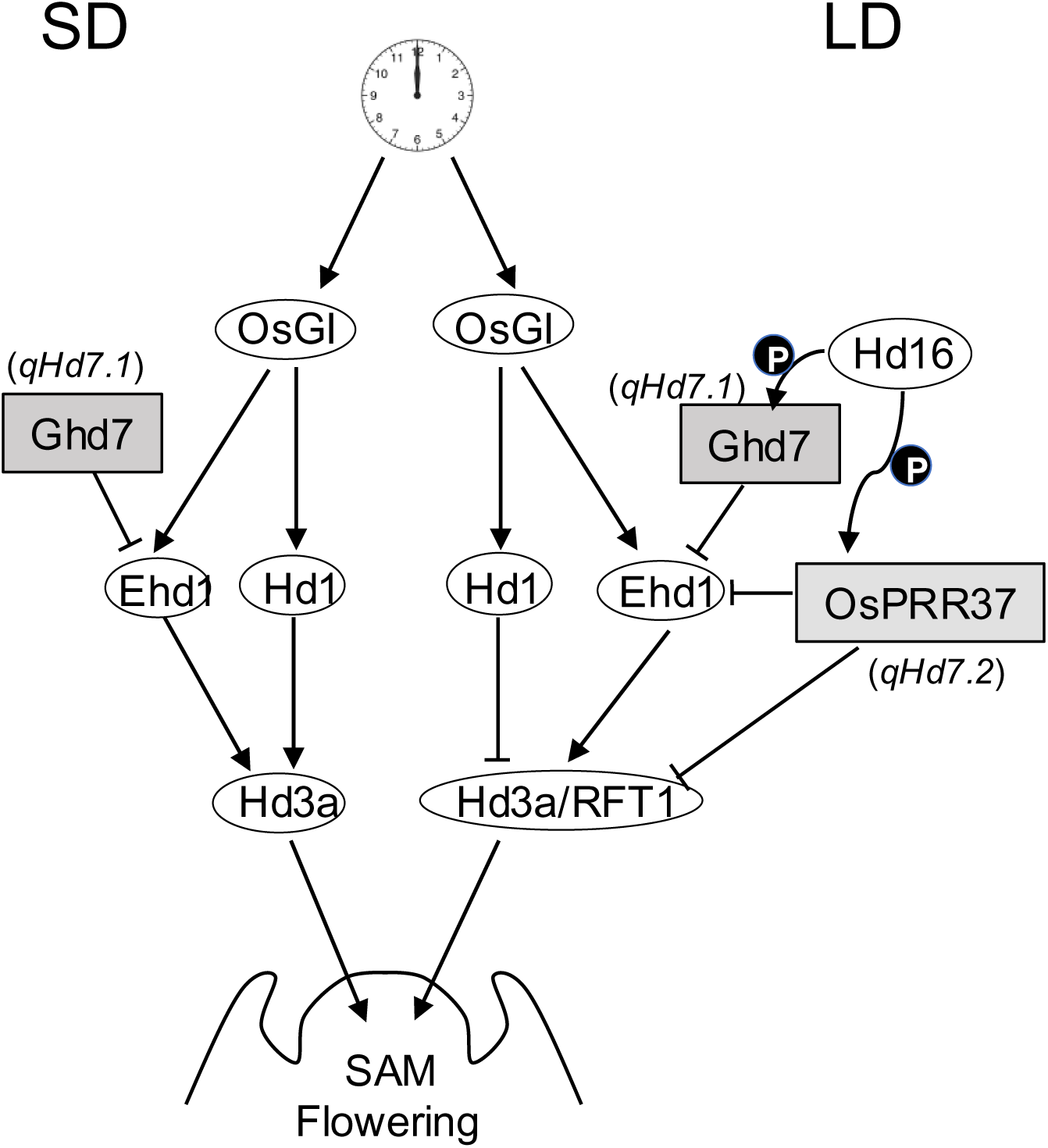
Schematic model of the gene regulatory network for the flowering pathway of rice under SD and LD conditions. Arrows and “T” sign indicates promotion and suppression, respectively. P indicates protein phosphorylation. Candidate causal genes for *qHd7.1* and *qHd7.2* are shown in gray rectangles. SD, short-day; LD, long-day; SAM, shoot apical meristem.

The DJ123 allele at *qHd7.1* caused early heading under both SD and LD conditions. Since *qHd7.1* is closely linked to the previously reported *Ghd7*, we will further examine evidence as to whether *qHd7.1* may be allelic to *Ghd7*. It has been reported that *Hd16* encodes a casein kinase I-like protein, and phosphorylation of Ghd7 by Hd16 increases the Ghd7 activity, which suppresses the expression of *Ehd1* and thereby delays flowering under LD conditions (Hori *et al*. 2013; Xue *et al*. 2008) (Fig. 7). The casein kinase is a member of serine/threonine selective enzymes that phosphorylate serine or threonine. Our sequencing data revealed that Ghd7 had four amino acid substitutions between IR64 and DJ123, including a change from a glycine to a serine residue at the position 136 in the IR64 allele, causing late heading. Position 136 is a predicted phosphorylation site of Ghd7 with serine being the binding site, suggesting that this phosphorylation site was lost from the DJ123 (and Nipponbare) allele (Figs. 6, 7; Supplemental Fig. 3). We hypothesize that the 136th serine is an important residue for phosphorylation that up-regulates Ghd7 activity to suppress *Ehd1* under SD and LD conditions in the IR64 allele. The DJ123 allele of Ghd7 lacks this potentially important regulatory site and therefore does not suppress *Ehd1*, leading to early heading (Fig. 7). In fact, our results show that the DJ123 allele of *qHd7.1* causes early heading under both SD and LD conditions (Fig. 3). Thus, while further fine-mapping would be needed to prove *qHd7.1* is allelic to *Ghd7*, observed allelic differences between the parents of the mapping population are consistent with the proposed mode of action of Ghd7.

### Adaptability exemplified by early heading and high yield potential under Madagascar conditions

Grain yield in rice is closely related to heading date. Early or late flowering in rice causes reduced grain production through insufficient growth of vegetative organs or reduced fertility as temperature decrease late in the season (Xu *et al*. 2014, Song *et al*. 2015). Genes and QTL related to flowering time often have pleiotropic effects on plant height and grain yield. For example, null mutations in *Ghd7* promote early flowering and reduce grain production and plant height under LD conditions (Xue *et al*. 2008). It is therefore crucial to consider yield penalties when embarking on selection for early heading.

In this study, the combined DJ123 alleles at QTLs *qHd7.1* and *qHd7.2* accelerated DTH by almost 10 days under LD conditions in Japan, and while there was no significant difference in average grain yield between the allelic groups, it appeared that earliest heading lines did have reduced grain yield (Fig. 4). This contrasted with the situation in Madagascar where only *qHd7.1* was effective in shortening the time to heading and where very early and high-yielding lines could be identified (Figs.4, 5). This would indicate that *qHd7.1* did not exhibit pleiotropic effects under SD conditions in Madagascar. Marker-assisted selection for the DJ123 allele at *qHd7.1* therefore appears a promising strategy to contribute to the demands of African farmers in terms of shortening of the rice cultivation period. Selected lines identified here furthermore indicate that this can be achieved while simultaneously selecting for increased yield potential. The development of short duration and high-yielding rice varieties will be a crucial contribution to developing more climate-resilient rice production systems for SSA.

## Author Contribution Statement

K.K. and M.W. conceived the hypotheses, designed the experiment, performed the analysis and interpretations, and wrote the manuscript. M.R. and H.N.R. performed the field experiments and data collections. K.H. performed DNA sequence analysis and provided critical feedback and helped shape the research. Y.U. provided editorial comments and corrected figures.

## Acknowledgments

Special thanks to Dr. C. Nathan Hancock from the University of South Carolina Aiken and Dr. Toshiyuki Takai from Japan International Research Center for Agricultural Sciences (JIRCAS) for editing the manuscript. This research was supported by the Science and Technology Research Partnership for Sustainable Development (SATREPS), Japan Science and the Technology Agency (JST)/Japan International Cooperation Agency (JICA) (Grant No. JPMJSA1608). Matthias Wissuwa has been partly funded by the Deutsche Forschungsgemeinschaft (DFG, German Research Foundation) under Germany’s Excellence Strategy – EXC 2070 – 390732324.

## Supplementary Files

**Supplemental Fig. 1 Photoperiods in Japan and Madagascar**

**Supplemental Fig. 2 Scatter plot analyses comparing heading date and agronomical traits in the RIL population**

**Supplemental Fig. 3 Predicted protein sequences of GHD7 for three *Ghd7* alleles**

**Supplemental Fig. 4 Predicted protein sequences of OsPRR37**

**Supplemental Table 1 Primer sequences for Ghd7 and OsPRR37 used in this study.**

## Notes

### Competing Interest Statement

The authors have declared no competing interest.

