## Supplementary material for "A DJ123 allele at the heading date quantitative trait locus *qHd7.1* promotes early heading without yield penalties under natural short-day and low fertility conditions in Madagascar": suppl Figs 1-4; suppl Table 1

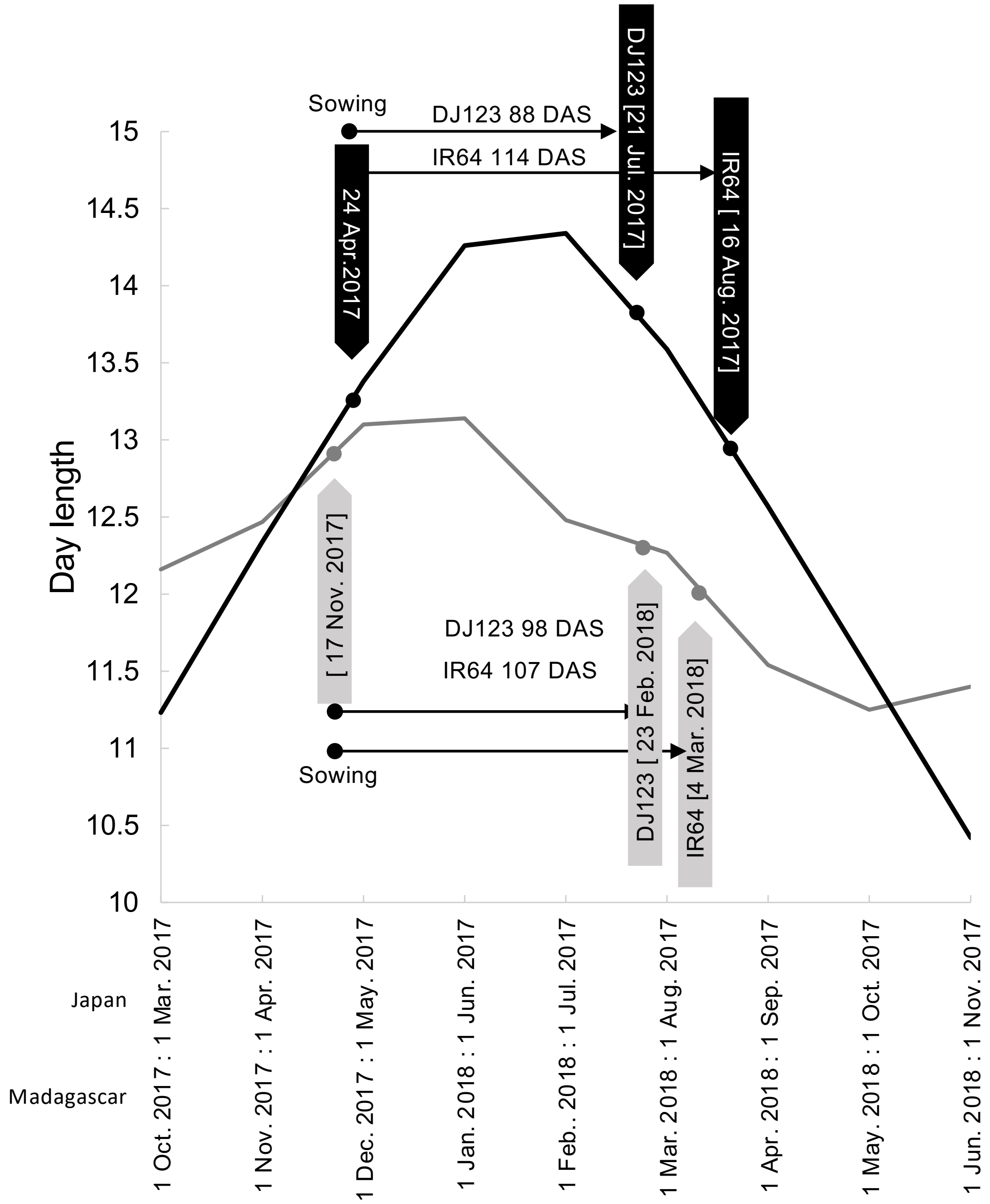


Supplemental Fig. 1 Photoperiods in Japan and Madagascar.
The field trials were conducted from April to September 2017 in Tsukuba, Japan, (36º05‘N, 140 º08’E) and from November 2017 to May 2018 in Ankazo, Madagascar, (19º40‘S, 46 º33’E). The black and gray lines indicate the day length of Japan LD and Madagascar SD conditions, respectively.

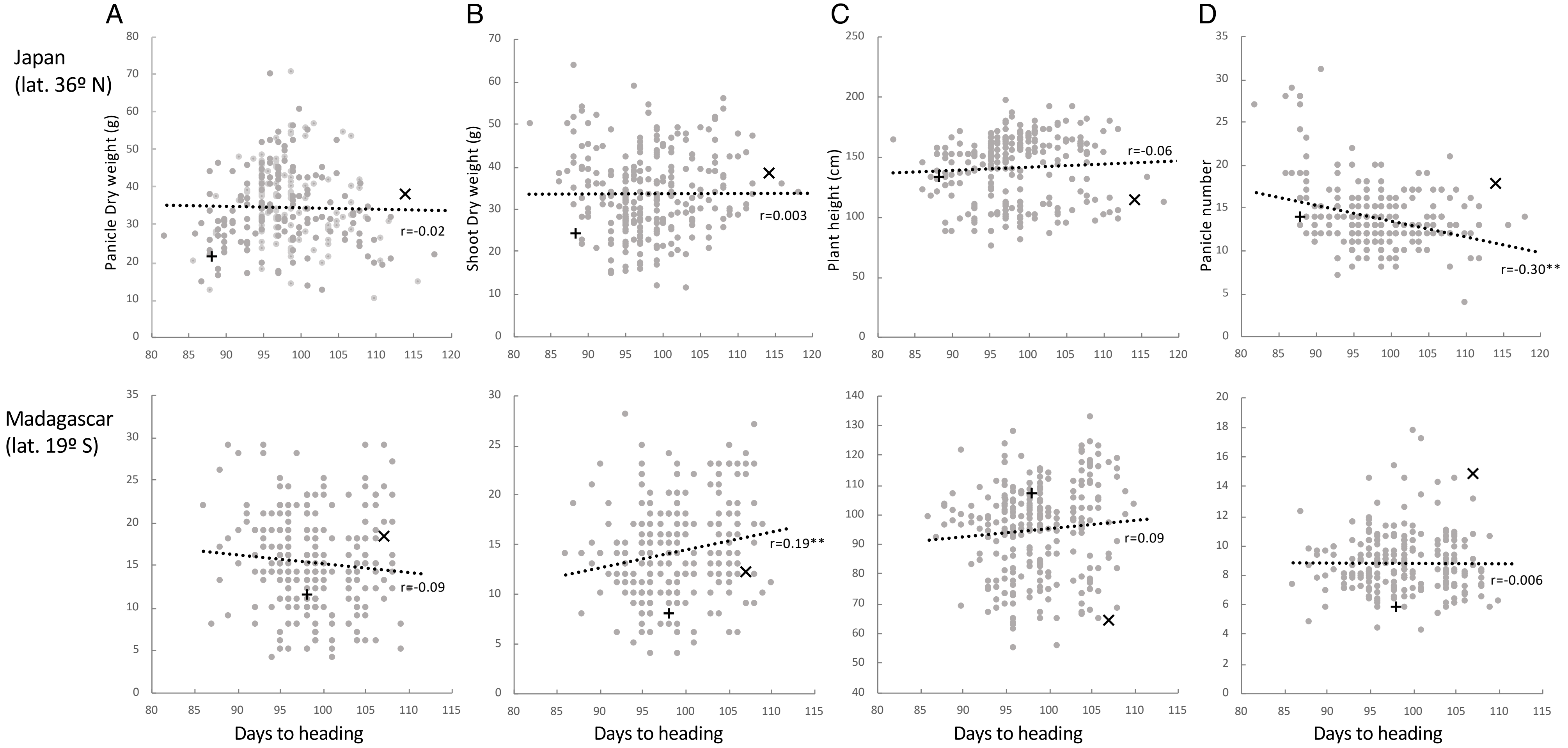


Supplemental Fig. 2 Scatter plot analyses comparing heading date and agronomical traits in the RIL population.
Data from Japan lat.36ºN under LD conditions, fertilizer application and Madagascar lat.19ºS under SD conditions, phosphorus-deficient conditions. Scatter plot comparing the panicle dry weight (A), shoot dry weight (B), plant height (C), panicle number (C) and days to heading, respectively. × and + indicate the average of parents IR64 and DJ123, respectively.

```
                                           *                            *
IR64 Ghd7 AA.seq      1 MSMGPAAGEGCGLCGADGGGCCSRHRHDDDGFPFVFPPSACQGIGAPAPPVHEFQFFGND   60
Ghd7 NB-AA.seq        1 MSMGPAAGEGCGLCGADGGGCCSRHRHDDDGFPFVFPPSACQGIGAPAPPVHEFQFFGND   60
DJ123 Ghd7 AA.seq     1 MSMGPAAGEGCGLCGADGGGCCSRHRHDDDGFPFVFPPSACQGIGAPAPPVHEFQFFGND   60

                                           *                            *
IR64 Ghd7 AA.seq     61 GGGDDGESVAWLFDDYPPPSPVAAAAGMHHRQPPYDGVVAPPSLFRRNTGAGGLTFDVSL  120
Ghd7 NB-AA.seq       61 GGGDDGESVAWLFDDYPPPSPVAAAAGMHHRQPPYDGVVAPPSLFRRNTGAGGLTFDVSL  120
DJ123 Ghd7 AA.seq    61 GGGDDGESVAWLFDDYPPPSPVAAAAGMHHRQPPYDGVVAPPSLFRRNTGAGGLTFDVSL  120

                                       *                    *    *
IR64 Ghd7 AA.seq    121 GGRPDLDAGLGLGGGSGRHAEAAASATIMSYCGSTFTDAASSMPKEMVAAMADVGESLNP  180
Ghd7 NB-AA.seq      121 GERPDLDAGLGLGGGGGRHAEAAASATIMSYCGSTFTDAASSMPKEMVAAMADDGESLNP  180
DJ123 Ghd7 AA.seq   121 GERPDLDAGLGLGGGGGRHAEAAASATIMSYCGSTFTDAASSMPKEMVAAMADDGESLNP  180

                                                         *
IR64 Ghd7 AA.seq    181 NTVVGAMVEREAKLMRYKEKRKKRCYEKQIRYASRKAYAEMRPRVRGRFAKEADQEAVAP  240
Ghd7 NB-AA.seq      181 NTVVGAMVEREAKLMRYKEKRKKRCYEKQIRYASRKAYAEMRPRVRGRFAKEPDQEAVAP  240
DJ123 Ghd7 AA.seq   181 NTVVGAMVEREAKLMRYKEKRKKRCYEKQIRYASRKAYAEMRPRVRGRFAKEPDQEAVAP  240

                         *     *
IR64 Ghd7 AA.seq    241 PSTYVDPSRLELGQWFR                                            257
Ghd7 NB-AA.seq      241 PSTYVDPSRLELGQWFR                                            257
DJ123 Ghd7 AA.seq   241 PSTYVDPSRLELGQWFR                                            257
```

Supplemental Fig. 3 Predicted protein sequences of GHD7 for three *Ghd7* alleles.
Gray shaded area represents the putative CCT domain (aa189-233). Asterisk indicates the phosphorylation site predicted by NetPhos 3.1 of serine/threonine kinase.

```
NB-AA.seq       1 MMGTAHHNQTAGSALGVGVGDANDAVPGAGGGGYSDPDGGPISGVQRPPQVCWERFIQKK    60
IR64-AA.seq     1 MMGTAHHNQTAGSALGVGVGDANDAVPGAGGGGYSDPDGGPTSGVQPPPQVCWERFIQKK    60
DJ123-AA.seq    1 MMGTAHHNQTAGSALGVGVGDANDAVPGAGGGGYSDPDGGPTSGVQPPPQVCWERFIQKK    60

NB-AA.seq      61 TIKVLLVDSDDSTRQVVSALLRHCMYEVIPAENGQQAWTYLEDMQNSIDLVLTEVVMPGV   120
IR64-AA.seq    61 TIKVLLVESDDSTRQVVSALLRHCMYEVIPAENGQQAWTYLEDMQNSIDLVLTEVVMPGV   120
DJ123-AA.seq   61 TIKVLLVESDDSTRQVVSALLRHCMYEVIPAENGQQAWTYLEDMQNSIDLVLTEVVMPGV   120

NB-AA.seq     121 SGISLLSRIMNHNICKNIPVIMMSSNDAMGTVFKCLSKGAVDFLVKPIRKNELKNLWQHV   180
IR64-AA.seq   121 SGISLLSRIMNHNICKNIPVIMMSSNDAMGTVFKCLSKGAVDFLVKPIRKNELKNLWQHV   180
DJ123-AA.seq  121 SGISLLSRIMNHNICKNIPVIMMSSNDAMGTVFKCLSKGAVDFLVKPIRKNELKNLWQHV   180

NB-AA.seq     181 WRRCHSSSGSGSESGIQTQKCAKSKSGDESNNNNGSNDDDDDDGVIMGLNARDGSDNGSG   240
IR64-AA.seq   181 WRRCHSSSGSGSESGIQTQKCAKSKSGDESNNNSGSNDDDDDDGVIMGLNARDGSDNGSG   240
DJ123-AA.seq  181 WRRCHSSSGSGSESGIQTQKCAKSKSGDESNNNSGSNDDDDDDGVIMGLNARDGSDNGSG   240

NB-AA.seq     241 TQAQSSWTKRAVEIDSPQAMSPDQLADPPDSTCAQVIHLKSDICSNRWLPCTSNKNSKKQ   300
IR64-AA.seq   241 TQAQSSWTKRAVEIDSPQAMSPDQLADPPDSTCAQVIHLKSDICSNRWLPCTSNKNSKKQ   300
DJ123-AA.seq  241 TQAQSSWTKRAVEIDSPQAMSPDQLADPPDSTCAQVIHLKSDICSNRWLPCTSNKNSKKQ   300

NB-AA.seq     301 KETNDDFKGKDLEIGSPRNLNTAYQSSPNERSIKPTDRRNEYPLQNNSKEAAMENLEESS   360
IR64-AA.seq   301 KETNDDFKGKDLEIGSPRNLNTAYQSSPNERSIKPTDRRNEYPLQNNSKEAAMENLEESS   360
DJ123-AA.seq  301 KETNDDFKGKDLEIGSPRNLNTAYQSSPNERSIKPTDRRNEYPLQNNSKEAAMENLEESS   360

NB-AA.seq     361 VRAADLIGSMAKNMDAQQAARAANAPNCSSKVPEGKDKNRDNIMPSLELSLKRSRSTGDG   420
IR64-AA.seq   361 VRAADLIGSMAKNMDAQQAARATNAPNCSSKVPEGKDKNRDNIMPSLELSLKRSRSTGDD   420
DJ123-AA.seq  361 VRAADLIGSMAKNMDAQQAARATNAPNCSSKVPEGKDKNRDNIMPSLELSLKRSRSTGDD   420

NB-AA.seq     421 ANAIQEEQRNVLRRSDLSAFTRYHTPVASNQGGTGFMGSCSLHDNSSEAMKTDSAYNMKS   480
IR64-AA.seq   421 ANAIQEEQRNVLRRSDLSAFTRYHTPVASNQGGTGFVGSCSPHDNISEAMKTDSAYNMKS   480
DJ123-AA.seq  421 ANAIQEEQRNVLRRSDLSAFTRYHTPVASNQGGTGFVGSCSPHDNISEAMKTDSAYNMKS   480

NB-AA.seq     481 NSDAAPIKQGSNGSSNNNDMGSTTKNVVTKPSTNKERVMSPSAVKANGHTSAFHPAQHWT   540
IR64-AA.seq   481 NSDAAPIKQGSNGSSNNNDMGSTTKNVVTKPSTNKERVMSPSAVKANGHTSAFHPAQHWT   540
DJ123-AA.seq  481 NSDAAPIKQGSNGSSNNNDMGSTTKNVVTKPSTNKERVMSPSAVKANGHTSAFHPAQHWT   540

NB-AA.seq     541 SPANTTGKEKTDEVANNAAKRAQPGEVQSNLVQHPRPILHYVHFDVSRENGGSGAPQCGS   600
IR64-AA.seq   541 SPANTTGKEKTDEVANNAAKRAQPGEVQSNLVQHPRPILHYVHFDVSRENGGSGAPQCGS   600
DJ123-AA.seq  541 SPANTTGKEKTDEVANNAAKRAQPGEVQSNLVQHPRPILHYVHFDVSRENGGSGAPQCGS   600

NB-AA.seq     601 SNVFDPPVEGHAANYGVNGSNSGSNNGSNGQNGSTTAVNAERPNMEIANGTINKSGPGGG   660
IR64-AA.seq   601 SNVFDPPVEGHAANYGVNGSNSGSNNGSNGQNGSTTAVNAERPNMEIANGTINKSGPGGG   660
DJ123-AA.seq  601 SNVFDPPVEGHAANYGVNGSNSGSNNGSNGQNGSTTAVNAERPNMEIANGTINKSGPGGG   660

NB-AA.seq     661 NGSGSGSGNDMYLKRFTQREHRVAAVIKFRQKRKERNFGKKVRYQSRKRLAEQRPRVRGQ   720
IR64-AA.seq   661 NGSGSGSGNDMYLKRFTQREHRVAAVIKFRQKRKERNFGKKVRYQSRKRLAEQRPRVRGQ   720
DJ123-AA.seq  661 NGSGSGSGNDMYLKRFTQREHRVAAVIKFRQKRKERNFGKKVRY----------------   704

NB-AA.seq     721 FVRQAVQDQQQQGGGREAAADR                                         742
IR64-AA.seq   721 FVRQAVQDQQQQGGGREAAADR                                         742
DJ123-AA.seq  704 ----------------------                                         704
```

Supplemental Fig. 4  Predicted protein sequences of OsPRR37.
Gray shaded area represents the putative CCT domain.

Supplemental Table 1. Primer sequences for *Ghd7* and *OsPRR37* used in this study.

| Primer name | Sequence (5'-3') |
|---|---|
| Ghd7-1U | ATGTCGATGGGACCAGCAG |
| Ghd7-1L | ACGCTCTCGCCGTCGTCGCCGC |
| Ghd7-2U | GCGGCGACGACGGCGAGAGCGT |
| Ghd7-2L | ATAGGTGGATGGCGGTGCGACA |
| Ghd7-3U | CTGTCGGACCGCCATCCACCTA |
| Ghd7-3L | TCCAAGAATTCCTATCTGAACCAT |
| OsPRR37-1U | CTCTTCGAGTCAGGCTGATGAT |
| OsPRR37-1L | AAGGAGGAAAGAAGAGACAGACC |
| OsPRR37-2U | CTTTGTTGGTCTTTGACTGAGTG |
| OsPRR37-2L | CTAAGAAGTCAACAGCGCCC |
| OsPRR37-3U | GGGCGCTGTTGACTTCTTAG |
| OsPRR37-3L | CATGGTGAACTCTTCCTATTTCC |
| OsPRR37-4U | AAGTGCATTTAAACTGGTTGCTC |
| OsPRR37-4L | AATTAAGTCAGCAGCTCGAACAC |
| OsPRR37-5U | TTATCAATCCTCTCCGAATGAGA |
| OsPRR37-5L | GTGTGTCCATTAGCCTTAACAGC |
| OsPRR37-6U | AACAATGACATGGGTTCCACTAC |
| OsPRR37-6L | AGCTAGTCGTTTGGGAGAAACTT |
| OsPRR37-7U | TCTCCCAAACGACTAGCTAAATG |
| OsPRR37-7L | ATCGCGTAGGTAGGTAGGTAGGT |
| Ghd7-pro1U | CTGGTGTGGAGGTGTGACTATC |
| Ghd7-pro1L | TGTGTGGGTGGTGGTATATTTTT |
| Ghd7-pro2U | TCGATGTTTTCTAACCGGAAATA |
| Ghd7-pro2L | AGAAAAACCTGCTTTTCAGCTCT |
| Ghd7-pro3U | AAGGGCCCCATATTATTGTTAAA |
| Ghd7-pro3L | AAGAAGAAGAAGACAGGGCAAGT |
| Ghd7-pro4U | TATGTTTCTCACACGGGATTTTT |
| Ghd7-pro4L | GTTGCCGAAGAACTGGAACTC |
